# MD Biophysics Photobiomodulation Plasma (PPT)/Very Small Embryonic like (VSEL) Antibody Marker Trend Analysis

**DOI:** 10.64898/2026.03.29.715134

**Authors:** Dawn DeSylvia, Ian Mitchell

## Abstract

**Background:** Photobiomodulation (PBM) therapy has demonstrated therapeutic potential in promoting cellular repair, modulating inflammation, and enhancing mitochondrial function. Platelet-rich plasma (PRP) is widely used in regenerative medicine due to its concentration of growth factors and cytokines. Very small embryonic-like stem cells (VSELs), a rare population of pluripotent stem cells present in adult tissues, have emerged as a potential contributor to tissue regeneration. While PBM and PRP are used in combination, how VSELs or Multi-lineage stress enduring (MUSE) cells are at play, and the biological mechanisms underlying their synergistic effects remain incompletely characterized.

**Objective:** This exploratory pilot study aimed to evaluate whether application of the MD Biophysics laser to autologous PRP is associated with measurable changes in VSEL-related antibody marker expression, and to identify directional trends to inform future controlled studies.

**Methods:** PRP samples were collected from participants across seven test dates (July 2024 to February 2025), yielding 18 participant-session datasets. Samples were analyzed before (Pre) and after (Post) laser application using flow cytometry conducted at a UCLA Flow Cytometry Laboratory. Four VSEL-associated antibody markers were assessed: CD45^−^CD34^+^, CXCR4^+^, CD133^+^, and SSEA-4^+^. Analyses were descriptive and focused on paired differences and directional trends due to the exploratory design and absence of a control group.

**Results:** Three of four VSEL-associated markers (CXCR4^+^, CD133^+^, and SSEA-4^+^) demonstrated a group-level increase in median paired differences following laser application. Directional increases were observed in 12/18 sessions for CXCR4^+^, 10/18 for CD133^+^, and 9/18 for SSEA-4^+^. CD45^−^CD34^+^ showed a near-equal distribution of increases and decreases. Ki-67 positivity indicated the presence of viable, proliferative cells. While no findings reached statistical significance due to limited sample size, consistent directional trends were observed across multiple markers.

**Conclusion:** Application of PBM to autologous PRP was associated with directional increases in multiple VSEL-associated antibody markers, suggesting a potential role for stem cell activation or mobilization in the mechanism of action. Although preliminary and not statistically powered, these findings provide hypothesis-generating evidence supporting further investigation. The observed trends informed iterative protocol refinement and establish a foundation for future controlled, adequately powered studies to evaluate clinical efficacy and underlying biological mechanisms.

## INTRODUCTION AND BACKGROUND

### Photobiomodulation Therapy

Photobiomodulation (PBM) therapy, also referred to as low-level light therapy or red-light therapy, involves the application of specific wavelengths of light to biological tissue to promote cellular repair and regeneration. The therapeutic potential of light has been recognized since antiquity, with early documentation of chromotherapy (color light therapy) and heliotherapy (sunlight therapy) in ancient Egyptian and Greek traditions [1, 2]. In the late nineteenth and early twentieth centuries, Dinshah Ghadiali developed early light-based instruments and proposed that specific spectral properties could influence physiological states [1, 2].

Contemporary scientific interest in PBM expanded following the work of biophysicist Fritz-Albert Popp in the 1970s, who identified that living cells emit coherent photons, termed biophotons, suggesting a role for light in intracellular communication [3, 4]. Structural components including fascia, collagen, and mitochondria are thought to facilitate biophoton transmission through mechanisms analogous to fiber optics, with implications for cellular coordination and disease states [3-7]. Professor Micheal Hamblin of Harvard has furthered the understanding and mechanisms of action of PBM in its ability to stimulate healing, relieve pain, and reduce inflammation [8].

More than 3000 peer-reviewed publications have examined the biological effects of red-light therapy [7]. In 2013, research at the University of Wisconsin demonstrated that near-infrared light activates mitochondrial cytochrome C oxidase, stimulating cellular repair processes, with experimental models showing efficacy in treating demyelinating disease and reversing retinal degeneration [8-12].

### Platelet-rich Plasma

Platelet-rich plasma (PRP) is an autologous preparation derived from centrifugation of whole blood, yielding a plasma fraction concentrated in platelets, growth factors, and cytokines involved in tissue repair and immune modulation. PRP has been widely evaluated as a vehicle for regenerative treatment, with documented applications in orthopedics, wound healing, and dermatology [13,14]. The combination of PBM and PRP has been applied clinically, with practitioners reporting synergistic effects; however, the underlying mechanisms remain incompletely characterized and there is lack formal peer-reviewed investigation.

### Very Small Embryonic-Like Stem Cells

Very Small Embryonic-Like stem cells (VSEL) are a rare population of pluripotent stem cells identified in adult bone marrow and peripheral tissues [15-19]. Initially described more than 2 decades ago, their presence has since been confirmed by more than 20 independent laboratories, with evidence of pluripotency marker expression and the capacity to differentiate across germ layers [15-21].

Compared with mesenchymal stem cells (MSC), VSEL exhibit several properties relevant to regenerative medicine. MSC are multipotent and undergo telomeric shortening commensurate with donor age.

By contrast, VSEL demonstrate negligible telomeric degradation, suggesting retention of a more primitive epigenetic profile. Their small diameter (5 to 7 microns, compared with 10 to 12 microns for Muse cells and 15 to 20 microns for MSC) may permit passage across the blood-brain barrier and other tissue boundaries that restrict larger cell populations. The VSEL surface phenotype is characterized by SSEA-4 expression among other pluripotency markers, distinguishing them from Muse cells, which are identified by SSEA-3 expression.

### Previous Work on VSEL Activation

Previous investigations have explored whether physical and energetic modalities can stimulate VSEL proliferation. In the early 2000s Dr John Wong, an immunologist and researcher, demonstrated that the application of specific frequencies and modalities to PRP promoted expansion of nonembryonic totipotent blastomere-like stem cells [21]. A decade later Dr Todd Ovokitys reported that VSEL exposed to laser light demonstrated increased proliferation relative to controls [20]. Additional investigators have examined VSEL activation through hormetic stressors including hypothermic conditions, mechanostimulation, and acoustic stimulation. In 2020, Ian Mitchell began development on laser quantization of cells employing 2 adjustable frequencies with pulse width modulation (PWM), enabling depth-specific energy delivery and selective activation of cellular layers, which enhanced therapeutic outcomes and expanded the applicability of laser-based plasma activation. Subsequently, he transitioned to a multiple channel entrainment system affording the ability to reach more tissues with specificity and deliver a more biologically appropriate Joules per cubic centimeter (J/cm3) of photonic energy to those regions. The development of this followed a formulaic approach incorporating the variable conditions of time, frequency, pulse width, and tissue density. Culminating in the development of the following formula which is the backbone of the MD Biophysics System:

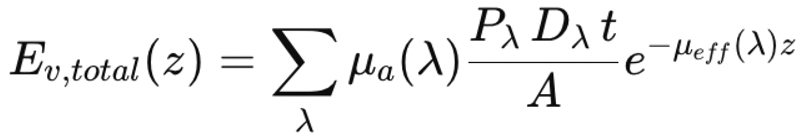

Notwithstanding these findings, a literature review and consultation with scientists at the UCLA Flow Cytometry Laboratory identified no peer-reviewed publication providing definitive evidence that light or frequency directly activates or expands VSEL or Muse cells. Furthermore, previous studies in this domain have generally employed a single VSEL antibody marker (CD45+/CD34-), and none have confirmed the presence of live, actively dividing cells. The present investigation addresses both limitations by employing a 4-marker antibody panel and incorporating Ki-67 as an indicator of active cell division.

### Study Rationale and Hypothesis

In May 2024, Dr Dawn DeSylvia (The Center for Whole Health, Agoura Hills, CA USA) and Ian Mitchell (Wizard Sciences, Bartlesville, OK USA) undertook an investigation to characterize the mechanism underlying clinical effects attributed to combined PBM and PRP treatment. Given the existing literature on PBM and VSEL biology, the central question was whether application of the MD Biophysics Laser to autologous PRP produces measurable changes in VSEL-associated antibody markers. In June 2024, the UCLA Flow Cytometry Laboratory (Los Angeles, CA USA) was contracted to quantify the percentage of VSEL antibody marker-positive cells in PRP specimens collected before and after laser application.

The MD Biophysics Laser is designed to modulate frequency for specific therapeutic applications and adjust depth of tissue penetration. For this investigation, a standardized wellness frequency set was applied.

The primary hypothesis was application of the MD Biophysics Laser to autologous PRP, using a defined collection and treatment protocol incorporating thermal, mechanical, and acoustic modalities, produces an increase in VSEL antibody marker expression.

The stated goals of the research were to characterize the mechanism of action of PBM applied to PRP and determine whether VSEL expansion constitutes a component of that mechanism; and) to identify the protocol and collection kit combination most likely to produce reproducible results, to support the development of a formalized clinical study.

This initial phase was designed as an exploratory pilot to identify directional trends rather than establish statistical significance. Protocol and collection kit selection proceeded iteratively across 7 test dates based on observed trends in VSEL antibody marker percentages after laser application.

### Recruitment

Patients of Dr. Dawn DeSylvia were invited to participate anonymously on a voluntary basis. Informed consent was obtained from all participants. Compensation consisted of a nutrient IV infusion. Participants donated peripheral blood for PRP preparation. The cohort included 18 individuals ranging in age from 32 to 90 years, with 11 women and 7 men participating.

## Methods

Data were collected across 7 study dates between July 2024 and February 2025. At each session, PRP was prepared from participant blood and the percentage of VSEL antibody marker-positive cells in PRP at baseline (before treatment) and after laser application, was assessed by flow cytometry analysis at a flow cytometry laboratory at the University of California. A range of PRP collection kits, laser devices, frequencies, and supplementary hormetic modalities (thermal, mechanical, acoustic) were evaluated across study dates. The final protocol incorporating the MD Biophysics Laser with an optimized frequency set was identified based on observed trends. Flow cytometry analysis quantified 4 VSEL-associated antibody markers: CD45-CD34+, CXCR4+, CD133+, and SSEA-4+. Ki-67 was added in the final study round to confirm the presence of actively dividing cells.

### Summary of Findings

Initial analysis suggested that the MD Biophysics Laser produced an increased number of positive antibody markers associated with VSEL presence, including in the context of Ki-67 positivity, confirming that live, actively dividing cells were detected. However, given the small sample size, statistical significance cannot be established.

## OVERVIEW

This paper presents a trend analysis of antibody marker expression measured by flow cytometry before and after laser application. Data were collected across 7 test days (July 2024 to February 2025), yielding 18 unique participant-session records. Four VSEL antibody markers were assessed: CD45^−^CD34^+^, CXCR4^+^, CD133^+^, and SSEA-4^+^. Time points are defined as Pre (base measurement for that participant within their test session) and Post (Laser 1 measurement for that participant within their test session). All analyses are descriptive and hypothesis-generating given the sample size and absence of a control group.

### Cytometry Database

VSEL marker values by session by participant are listed in Table 1

**Table 1:**
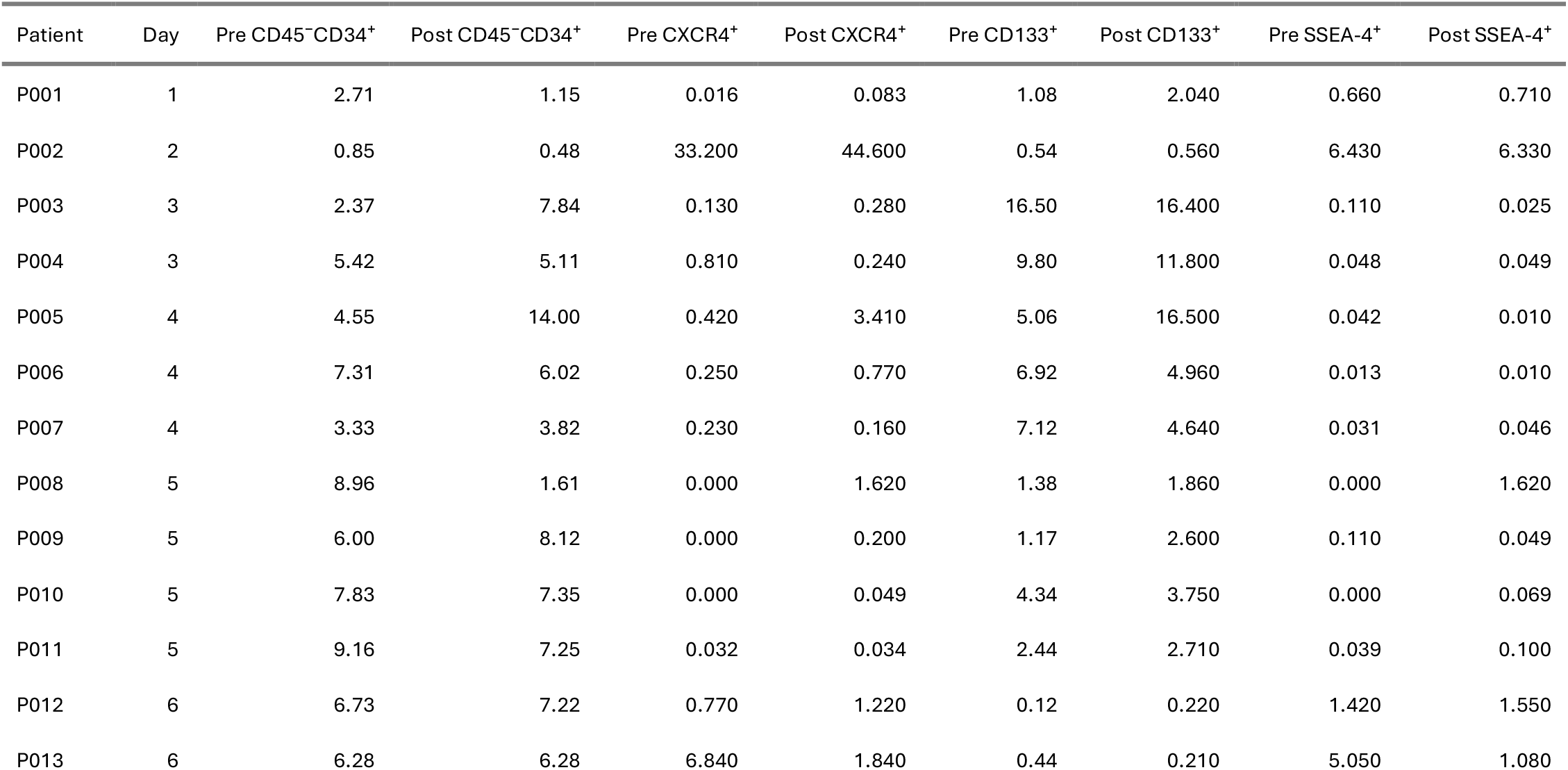
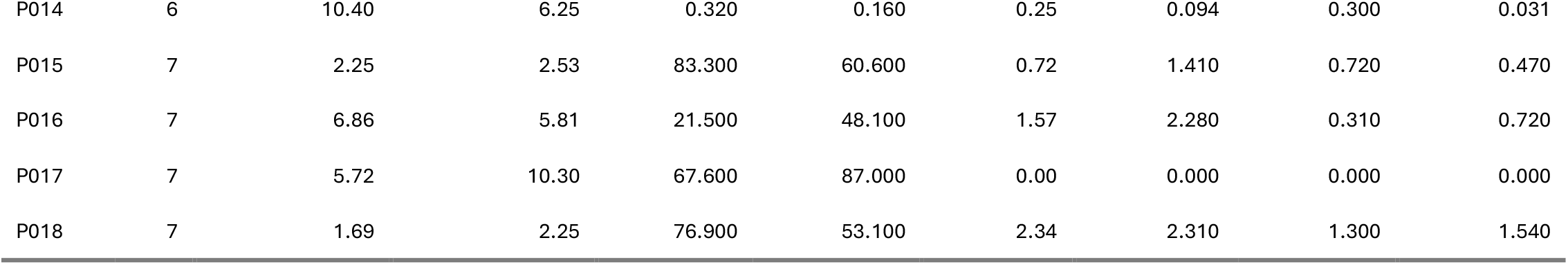
VSEL Marker Values by Participant-Session (Pre = Base; Post = Laser 1 within session)

### Descriptive Statistics

Marker values by time point are provided in Table 2.

**Table 2:**
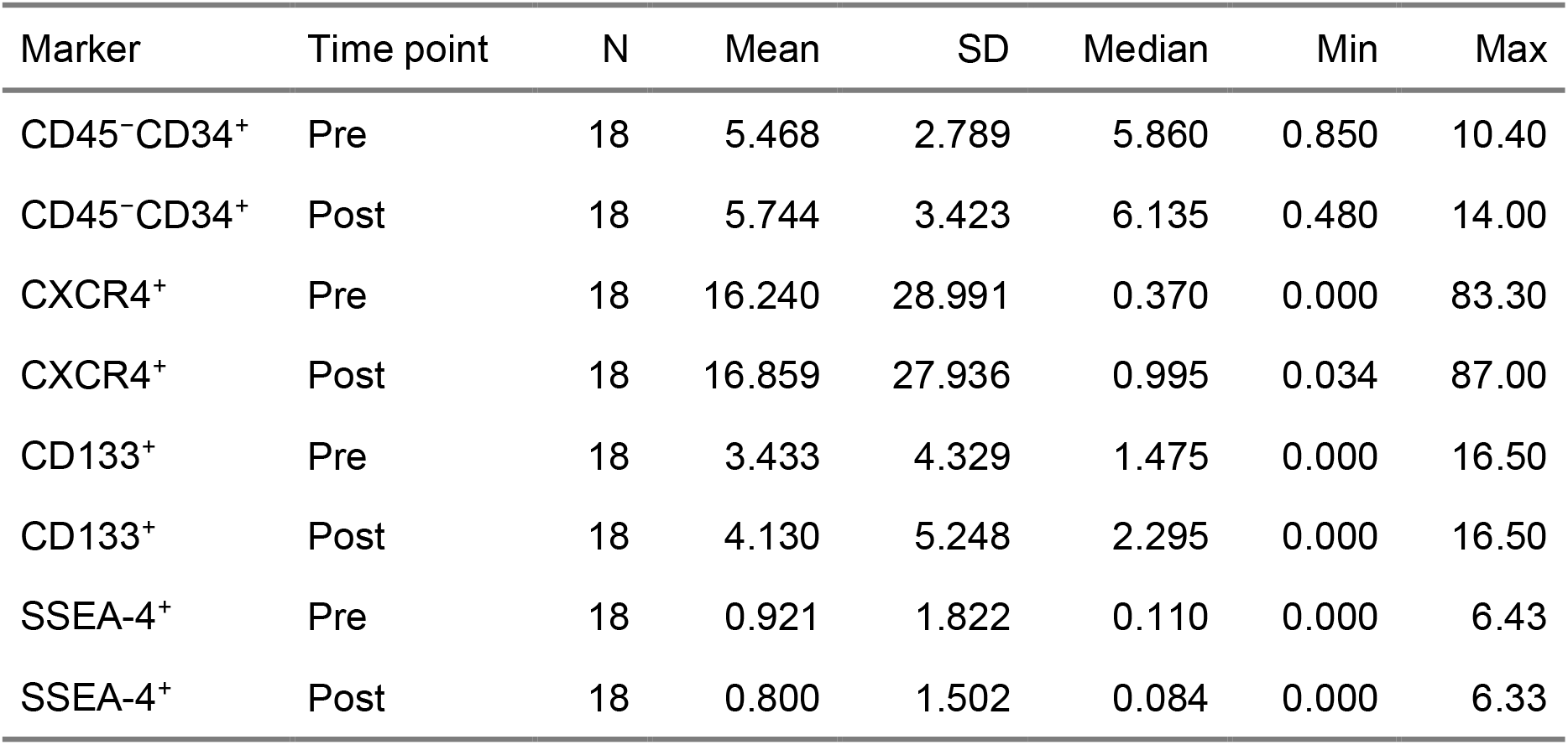
Descriptive Statistics: Marker Values by Time point.

### Visualizations

#### Grouped Bar Chart by Marker

Bars represent group means with standard error of the mean. Individual participant-session values are shown as overlaid points. Figure1 illustrates the mean Pre versus Post VSEL antibody marker values by time points.

**Figure 1:**
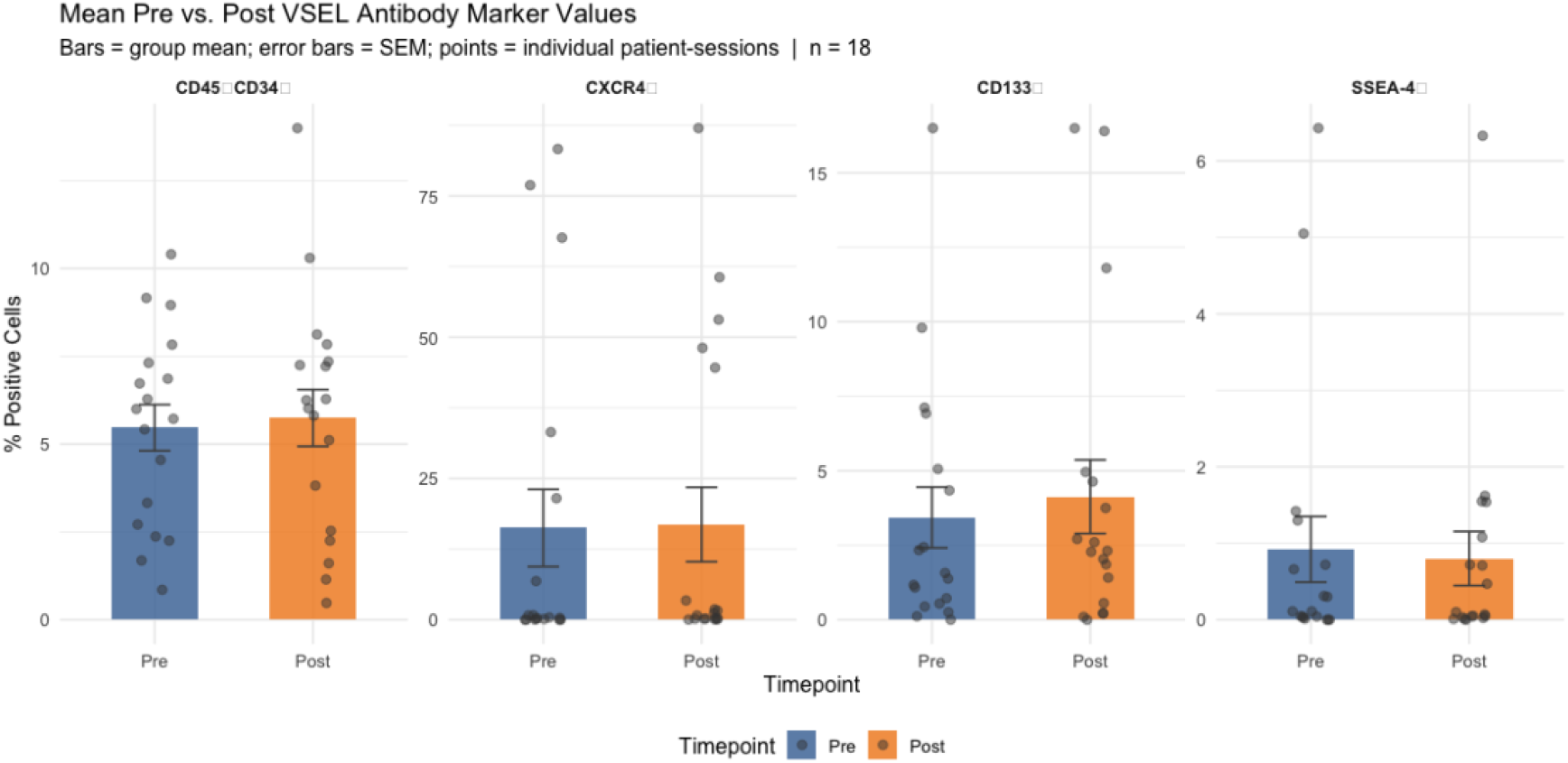
Mean Pre *versus* Post VSEL antibody marker values by time point. Bars represent group means; error bars represent standard error of the mean. Points show individual patient-session values. n = 18.

#### Paired Line Plot by Marker

Figure 2 illustrates Pre versus Post VSEL antibody markers. Each line represents one participant-session, connecting Pre (Base) and Post (Laser 1) values. Green lines indicate an increase from Pre to Post; red lines indicate a decrease.

**Figure 2:**
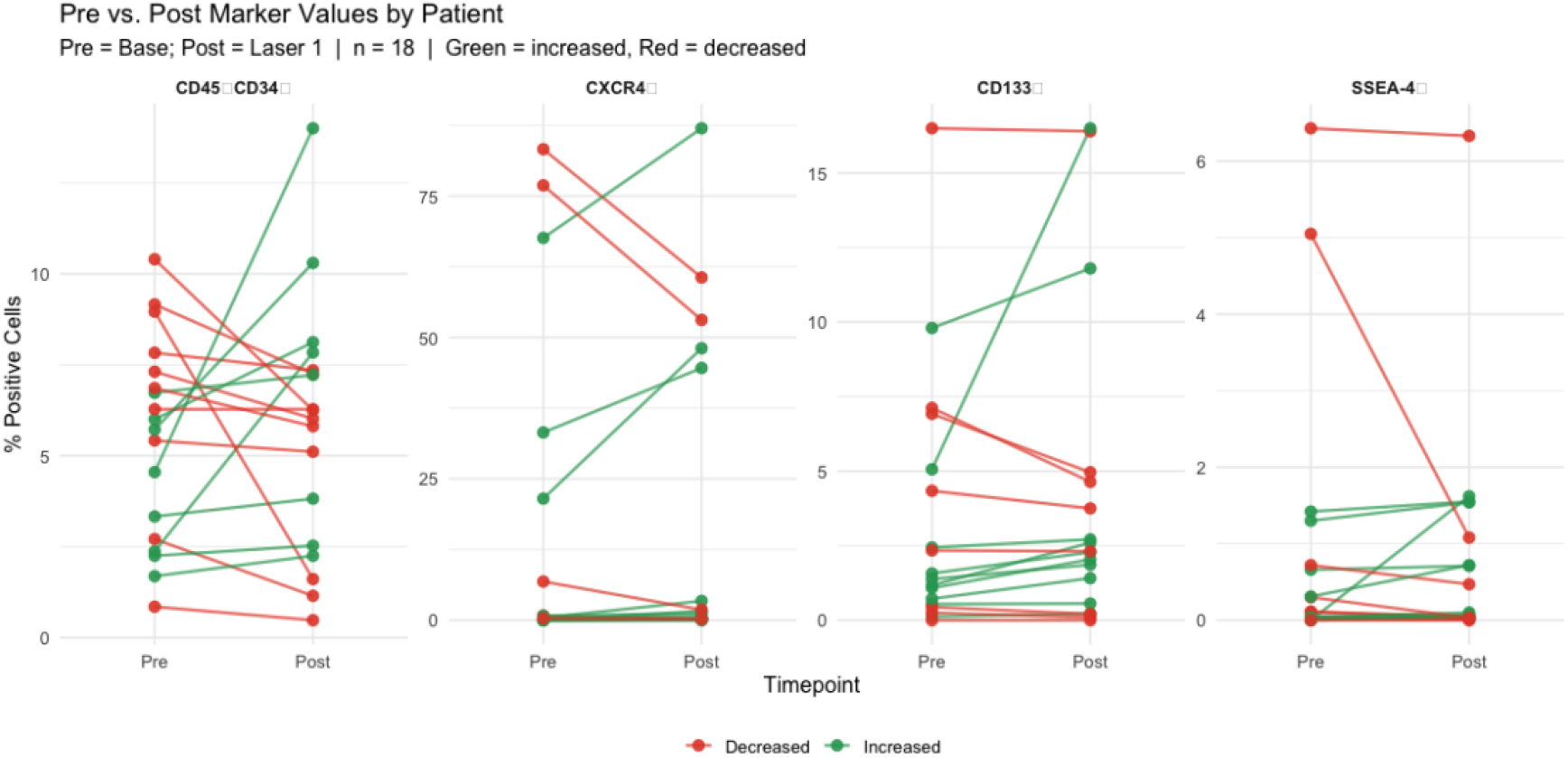
Pre *versus*. Post VSEL antibody marker values by patient-session. Each line connects one patient’s Pre (Base) and Post (Laser 1) measurements. Green = increased; red = decreased. n = 18.

#### Box Plots with Individual Data Points

The distribution of VSEL antibody marker values before and after laser application is illustrated in Figure 3.

**Figure 3:**
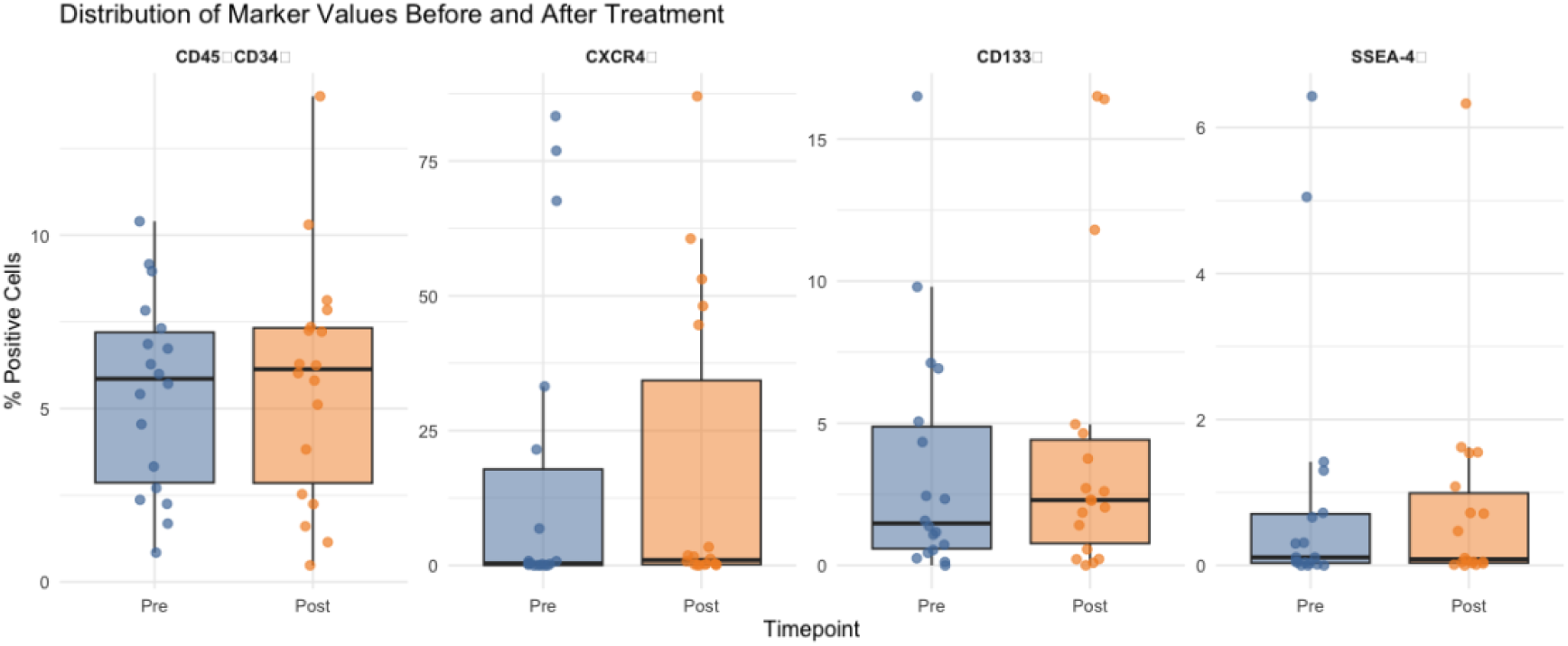
Distribution of VSEL antibody marker values before and after laser application. Boxes show the interquartile range; points show individual *participant*-session values.

#### Percent Change from Baseline

Data are provided illustrating the mean percent change in VESL antibody marker values from Pre and Post evaluations (Figure 4).

**Figure 4:**
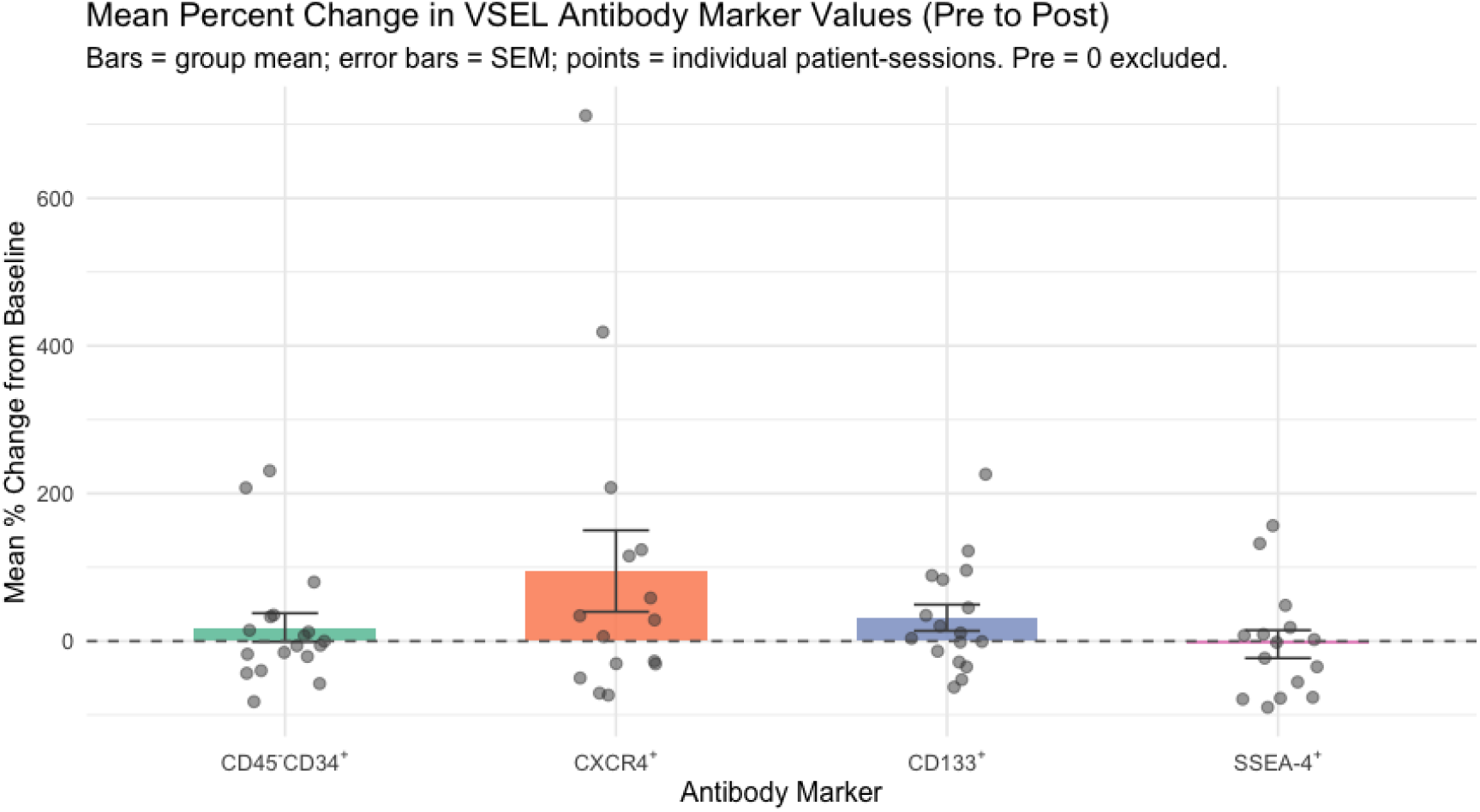
Mean percent change in VSEL antibody marker values from Pre to Post. Bars represent group means; error bars represent standard error of the mean. Points show individual patient-session values. Patient-sessions with a Pre value of 0 are excluded (percent change undefined).

## STATISTICAL TESTING

### Direction of Change

For each marker, Table 3 shows how many participants increased versus decreased from Pre to Post, and the overall group direction based on the median change.

**Table 3:** Direction of Change: Pre to Post by Marker.

| Marker | N Increased | N Decreased | N No Change | Median Change | Group Direction |
| --- | --- | --- | --- | --- | --- |
| CD45<sup>-</sup>CD34<sup>+</sup> | 8 | 9 | 1 | -0.155 | ↓ Decreased |
| CXCR4<sup>+</sup> | 12 | 6 | 0 | 0.109 | ↑ Increased |
| CD133<sup>+</sup> | 10 | 7 | 1 | 0.060 | ↑ Increased |
| SSEA-4<sup>+</sup> | 9 | 8 | 1 | 0.001 | ↑ Increased |

### Wilcoxon Signed-Rank Test (Paired, Non-Parametric

Given the small sample size, a non-parametric paired test was used. Results should be interpreted with caution given n = 18 (Table 4).

**Table 4:** Wilcoxon Signed-Rank Test Results (Paired Pre versus Post)

| Marker | W Statistic | P-value | Significance |
| --- | --- | --- | --- |
| CD45<sup>-</sup>CD34<sup>+</sup> | 77 | 1.0000 | ns |
| CXCR4<sup>+</sup> | 105 | 0.4080 | ns |
| CD133<sup>+</sup> | 97 | 0.3438 | ns |
| SSEA-4<sup>+</sup> | 78 | 0.9622 | ns |

\* p<0.05; . p<0.10; ns = not significant. Small sample size limits power.

### Effect Size (Cohen’s d, Paired)

Cohen’s d for paired samples was calculated as the mean of paired differences divided by the standard deviation of those differences. Conventional benchmarks for interpreting magnitude are shown in Table 5 and Table 6 [22].

**Table 5:** Cohen’s d Interpretation Benchmarks [22].

| Cohen's d | Magnitude | Interpretation |
| --- | --- | --- |
| < 0.20 | Negligible | Effect is smaller than typical measurement noise |
| 0.20 - 0.49 | Small | Subtle effect, may lack practical significance |
| 0.50 - 0.79 | Medium | Moderate effect, practically meaningful |
| >= 0.80 | Large | Substantial effect, clearly detectable |

**Table 6:** Paired Effect Sizes (Cohen’s d)

| Marker | Cohen's d | Magnitude |
| --- | --- | --- |
| CD45<sup>-</sup>CD34<sup>+</sup> | 0.076 | Negligible |
| CXCR4<sup>+</sup> | 0.053 | Negligible |
| CD133<sup>+</sup> | 0.242 | Small |
| SSEA-4<sup>+</sup> | -0.116 | Negligible |

## SUMMARY AND INTERPRETATION

### Summary Table

The data are summarized in Table 7.

**Table 7:** Summary: Pre/Post Values, Direction, Statistical Test, and Effect Size.

| Marker | Pre Mean | Pre SD | Post Mean | Post SD | N ↑ | N ↓ | Direction | P-value | Sig | Cohen's d | Effect Size |
| --- | --- | --- | --- | --- | --- | --- | --- | --- | --- | --- | --- |
| CD45<sup>-</sup>CD34<sup>+</sup> | 5.468 | 2.789 | 5.744 | 3.423 | 8 | 9 | ↓ Decreased | 1.0000 | ns | 0.076 | Negligible |
| CXCR4<sup>+</sup> | 16.240 | 28.991 | 16.859 | 27.936 | 12 | 6 | ↑ Increased | 0.4080 | ns | 0.053 | Negligible |
| CD133<sup>+</sup> | 3.433 | 4.329 | 4.130 | 5.248 | 10 | 7 | ↑ Increased | 0.3438 | ns | 0.242 | Small |
| SSEA-4<sup>+</sup> | 0.921 | 1.822 | 0.800 | 1.502 | 9 | 8 | ↑ Increased | 0.9622 | ns | -0.116 | Negligible |

### Interpretation

#### Overview

Across the 4 VSEL-associated antibody markers evaluated, 3 of 4 showed a group-level increase in the median paired difference from Pre to Post. No marker reached conventional statistical significance at this sample size (n = 18), which is expected given the exploratory design and limited power. Absence of significance does not indicate absence of effect; the observed effect sizes and directional patterns provide meaningful signal for hypothesis refinement and future study design. Substantial interpatient variability was present across all markers, reflecting the heterogeneity inherent to autologous PRP protocols and differences in individual baseline VSEL mobilization status.

#### CD45^-^CD34^+^

CD45^−^CD34^+^ is the most widely used VSEL surface marker in the literature, reflecting a hematopoietic progenitor phenotype. In this dataset, 8 of 18 participant-sessions showed an increase after laser application and 9 showed a decrease, with a median paired change of -0.155% (group direction: Decreased). The Wilcoxon signed-rank test was nonsignificant (p = 1), and the paired effect size was negligible (Cohen’s d = 0.076). The near-equal split between responders and nonresponders suggests that this marker may be sensitive to protocol variability, collection kit differences, or individual baseline differences in circulating progenitor cells. The presence of a subset of clear responders warrants further investigation with a standardized protocol.

#### CXCR4^+^

CXCR4^+^ is a chemokine receptor associated with stem cell homing and mobilization, and its expression on VSEL is thought to facilitate tissue-directed migration. In this dataset, 12 of 18 participant-sessions showed an increase and 6 showed a decrease (median change: 0.109%; group direction: ↑ Increased). The Wilcoxon test was nonsignificant (p = 0.408), with a negligible effect size (Cohen’s d = 0.053). This marker showed the widest absolute variability in the dataset, with several Day 7 participant-sessions exhibiting very high baseline CXCR4 expression (>60%) relative to earlier test days. This heterogeneity substantially inflates variance and reduces statistical power. The majority-increase pattern, despite high variability, is consistent with a mobilization effect of laser application on CXCR4-expressing progenitor cells and merits replication in a controlled cohort with matched baselines.

#### CD133^+^

CD133^+^ marks a population of primitive progenitor cells with pluripotent characteristics and is frequently used alongside CD45^−^CD34^+^ to characterize VSEL-enriched fractions. In this dataset, 10 of 18 participant-sessions showed an increase and 7 showed a decrease (median change: 0.06%; group direction: Increased). The Wilcoxon test was nonsignificant (p = 0.3438), with a small effect size (Cohen’s d = 0.242). CD133^+^ showed the most consistent directional pattern of the 4 markers, with most of participant-sessions trending toward increase. Although 1 participant-session contributed a notably large increase that influences the mean, the directional majority is consistent across a broad range of baseline values and collection days, strengthening the plausibility of a treatment-associated effect on this marker.

#### SSEA-4^+^

SSEA-4 (Stage-Specific Embryonic Antigen-4) is a glycan epitope expressed on pluripotent stem cells and is one of the markers distinguishing VSEL from Muse cells in the current classification framework. In this dataset, 9 of 18 participant-sessions showed an increase and 8 showed a decrease (median change: 0.001%; group direction: ↑ Increased). The Wilcoxon test was nonsignificant (p = 0.96Eff), with a negligible effect size (Cohen’s d = -0.116). SSEA-4 exhibited the smallest absolute values and the most balanced response distribution, suggesting either that this marker is less responsive to the treatment modality evaluated, that the detection threshold and assay sensitivity may limit signal resolution at low abundance, or that the relatively small SSEA-4-positive fraction is subject to disproportionate noise from minor cell count fluctuations.

#### Interpreting Non-Significant Findings in a Pilot Context

The absence of statistical significance across all 4 markers should not be interpreted as evidence of no effect. With n = 18, a Wilcoxon signed-rank test achieves limited statistical power, particularly in the presence of the high interpatient variability observed here. To detect a medium effect (Cohen’s d = 0.5) with 80% power at the Bonferroni-corrected 2-sided alpha = 0.0125 in a paired design, a substantially larger sample than the current n = 18 is required, as detailed in a later section (Future Directions). The observed effect sizes (ranging from 0.053 to 0.242 in absolute value) indicate that at least some markers are exhibiting detectable signal relative to within-subject variability, and these estimates provide a data-driven basis for powering future confirmatory studies.

Several factors beyond sample size contribute to the null findings and should be addressed in subsequent work. Protocol heterogeneity across the 7 test dates, including variation in collection kits, laser devices, and procedural parameters, introduces systematic noise that would attenuate any true treatment signal. Additionally, without a concurrent control group, regression to the mean, diurnal variation in circulating stem cell populations, and measurement variability at the flow cytometry level cannot be excluded as alternative explanations for observed changes. Despite these limitations, the directional consistency seen in CD133^+^ and, to a lesser extent, CXCR4^+^ suggests that specific markers may be more tractable endpoints for future confirmatory trials. Prioritizing these markers in a standardized, controlled protocol design would maximize the probability of detecting a true treatment effect.

#### Directional Trends as the Basis for Protocol Refinement

A key question in this exploratory investigation was whether any consistent directional signal could be observed across participant-sessions, even in the absence of statistical significance. Across 3 of 4 markers, the answer is yes.

Across the 18 participant-sessions analyzed, CXCR4^+^ showed an increase in 12 of 18 participant-sessions after laser application, with 6 decreasing. CD133^+^ showed an increase in 10 of 18 participant-sessions, with 7 decreasing and 1 showing no change. SSEA-4^+^ showed an increase in 9 of 18 participant - sessions, with 8 decreasing and 1 showing no change. CD45^−^CD34^+^ was the only marker where decreases slightly outnumbered increases (9 decreased, 8 increased, 1 no change), though the split was close.

In an exploratory pilot of this nature, directional majority is a meaningful and appropriate basis for decision-making. Statistical significance requires sufficient sample size to detect a signal above noise.

Directional consistency across individuals, by contrast, can be observed even in small samples and provides early evidence that a treatment may be producing a real effect. These pilot data do not provide confirmation of an effect, but rather orient and inform the design and measurement of an adequately-powered study.

The directional patterns observed here, particularly for CXCR4^+^ and CD133^+^, provided the basis for iterative protocol refinement across the 7 study dates. Protocols, collection kits, and procedural parameters that appeared to generate more consistent directional responses were retained and prioritized. The outcome is not a statistically confirmed treatment effect, but a directionally informed protocol now positioned for formal evaluation in a powered, controlled study.

## ANALYTIC CAVEATS

In summary, the analysis must be considered in light of several analytical caveats.

- Small sample size (n = 18): Statistical power is limited. Observed trends should be considered hypothesis-generating rather than confirmatory.
- Single post-treatment time point: Post reflects the Laser 1 measurement only. Other treatment readings, where collected, are not included in this analysis.
- Zero baseline values: Several participants had Pre = 0 for CXCR4^+^ and SSEA-4^+^ (Days 5 and 7). Percent change calculations are undefined for these cases and were excluded from the percent change visualization.
- Uncontrolled variables: Without randomization or a control group, observed changes cannot be attributed solely to the device intervention.
- Multiple comparisons: Testing 4 markers simultaneously increases false-positive risk. Bonferroni-corrected threshold would be p < 0.0125.
- Non-parametric testing: Wilcoxon signed-rank tests were selected given small n and potential nonnormality of flow cytometry distributions.
- Recommended next steps: A pilot study with matched controls, larger n, and a prespecified primary endpoint would substantially strengthen inferential validity.

## CLINICAL SIGNIFICANCE

Although the precise mechanisms underlying the clinically meaningful outcomes observed following PPT treatments with the MD Biophysics laser remain to be fully elucidated, the present findings are consistent with a growing body of evidence in the field of photobiomodulation (PBM). Existing data from academic and clinical research institutions not only corroborate these outcomes but also provide insight into potential biological mechanisms and avenues for further investigation.

Early work from the University of Michigan (2013) demonstrated significant therapeutic benefits of phototherapy in both wound healing and ocular disease models. Specifically, phototherapy-treated wounds exhibited healing times reduced by more than 50% compared to untreated controls [9]. In ophthalmologic applications, improvements were observed in both inherited conditions, such as retinitis pigmentosa, and acquired diseases, including diabetic retinopathy [9]. These findings were attributed to the observation that chronic, nonhealing wounds or diseases are arrested in the inflammatory phase of repair, and that light therapy helps overcome this barrier, enabling the normal healing processes to resume.

More recent evidence published in the Journal of Biophotonics indicates that brief exposure (15 minutes) to 670 nm red light can reduce blood glucose levels and attenuate postprandial glucose spikes by approximately 27.7%, suggesting a potential adjunctive role for PBM in metabolic regulation and diabetes management [21].

Consistent with these reports, our clinical observations following MD Biophysics PPT treatments demonstrate measurable improvements across multiple physiological domains. These include enhanced visual function, improved glycemic control, accelerated resolution of structural and musculoskeletal injuries without the need for invasive interventions, and functional improvements in cardiovascular performance, including increases in ejection fraction of 10–15 percentage points in patients with congestive heart failure. Additionally, improvements have been observed in neurocognitive and neuropsychiatric conditions—including Parkinson’s disease, Alzheimer’s disease, obsessive-compulsive disorder, attention-deficit/hyperactivity disorder, autism spectrum conditions, anxiety, and depression—as well as sustained gains in color perception among individuals with congenital color vision deficiency.

A unifying mechanism increasingly supported across PBM research is the role of mitochondrial activation and subsequent increases in adenosine triphosphate (ATP) production. Mitochondria serve as central regulators of cellular metabolism, and their activity appears to be directly influenced by specific wavelengths of light. As reported in the Journal of Biophotonics, exposure to 670 nm red light enhances mitochondrial membrane potential and stimulates ATP synthesis, thereby increasing cellular energy availability [21]. This increase in bioenergetic capacity may enable dysfunctional or damaged cells to restore normal physiological function and initiate repair processes.

In addition to mitochondrial effects, the work of Michael Hamblin highlights several supportive mechanisms, including modulation of reactive oxygen species (ROS), activation of downstream gene expression pathways, and nitric oxide release [8]. These processes may collectively contribute to improved cellular signaling, vascular function, and tissue repair. The potential interaction of these mechanisms with very small embryonic-like stem cells (VSELs) and multilineage-differentiating stress-enduring (MUSE) cells—particularly in the context of platelet-rich plasma (PRP)—represents a promising area for future research.

While additional research is needed to fully define these mechanisms, the present findings reflect core principles within photobiology. As emphasized by Dr. Fritz-Albert Popp, light is not merely an external stimulus but a fundamental regulator of biological function, capable of initiating and modulating critical cellular processes.

> “We know today that man, essentially, is a being of light. And the modern science of photobiology … is presently proving this. In terms of healing, the implications are immense. We now know, for example, that quanta of light can initiate, or arrest, cascade-like reactions in the cells, and that genetic cellular damage can be virtually repaired, within hours, by faint beams of light. We are still on the threshold of fully understanding the complex relationship between light and life, but we can now say emphatically, that the function of our entire metabolism is dependent on light.” [3].

## FUTURE DIRECTIONS

In an exploratory pilot of this nature, directional majority is a meaningful and appropriate basis for decision-making. The results suggest that the treatment may offer positive improvement for patients and future studies may include several modifications. Recommendations for future research would include:

Three different lasers and 3 different PRP collection kits were evaluated across 7 different days. With adequate sample sizes, future studies could formally test the following aims.

Because all 4 VSEL antibody markers will be tested simultaneously in a formal study, a Bonferroni correction is applied to control the family-wise error rate. The adjusted significance threshold is alpha = 0.05 / 4 = 0.0125. All sample size calculations below use this corrected alpha. Selecting only the markers that showed directional increases in this pilot would constitute post-hoc endpoint selection and is not recommended.

The primary aim of future studies will focus on whether laser application produces a statistically significant expansion of the VSEL population (pre versus post, paired design). The sample size calculation in Table 8 assumes a paired t-test at 2-sided alpha = 0.0125 and 80% power. A medium effect size (Cohen’s d = 0.5) is used as a conservative planning estimate, consistent with the range of effects observed in this pilot.

**Table 8:** Sample Size Requirements for Future Studies.

| Aim | Test | Effect Size | Alpha | Power | N Required | Note |
| --- | --- | --- | --- | --- | --- | --- |
| Primary: Paired pre/post VSEL presence | Paired t-test | $d = 0.50$ (medium) | 0.0125 | 0.8 | 48 | 48 total participants |

Pilot Cohen’s d range: 0.05 to 0.24. Conservative medium effect used for planning. Alpha = 0.0125 reflects Bonferroni correction for simultaneous testing of 4 markers (0.05 / 4), controlling the family-wise error rate at 0.05.

In addition to further understanding a statistical significance of VSEL population before and after laser application; an increased understanding of the mechanism of action driving the significant clinical findings of PPT is of interest. The following are initial study directions are recommended: 1. The addition of SSEA 3-antibody marker to assess if the MD Biophysics laser would also show an increase in MUSE antibody markers. 2. Measurement of hormones and cytokines and growth factors, before and after MD biophysics laser application. 3. Provide before and after clinical outcomes alongside flow cytometry and other chemical mediator analysis.

## Acknowledgements

The authors gratefully acknowledge the biostatistical analysis and technical editing contributions provided by Dr Liana Bruce, *Bruce Analytic Consulting Services, LLC*. The authors gratefully acknowledge, Darcy DeSylvia, as research administrative assistant.

